# ZIFA: Dimensionality reduction for zero-inflated single cell gene expression analysis

**DOI:** 10.1101/019141

**Authors:** Emma Pierson, Christopher Yau

## Abstract

Single cell RNA-seq data allows insight into normal cellular function and diseases including cancer through the molecular characterisation of cellular state at the single-cell level. Dimensionality reduction of such high-dimensional datasets is essential for visualization and analysis, but single-cell RNA-seq data is challenging for classical dimensionality reduction methods because of the prevalence of dropout events leading to zero-inflated data. Here we develop a dimensionality reduction method, (Z)ero (I)nflated (F)actor (A)nalysis (ZIFA), which explicitly models the dropout characteristics, and show that it improves modelling accuracy on simulated and biological datasets.

## Introduction

Single cell RNA expression analysis (scRNA-seq) is revolutionizing whole-organism science [1, 2] allowing the unbiased identification of previously uncharacterized molecular heterogeneity at the cellular level. Statistical analysis of single cell gene expression profiles can highlight putative cellular subtypes, delineating subgroups of T-cells [3], lung cells [4] and myoblasts [5]. These subgroups can be clinically relevant: for example, individual brain tumors contain cells from multiple types of brain cancers, and greater tumor heterogeneity is associated with worse prognosis [6].

Despite the success of early single cell studies, the statistical tools that have been applied to date are largely generic, rarely taking into account the particular structural features of single cell expression data. In particular, single cell gene expression data contains an abundance of dropout events that lead to zero expression measurements. These dropout events may be the result of technical sampling effects (due to low transcript numbers) or real biology arising from stochastic transcriptional activity (Figure 1a). Previous work has been undertaken to account for dropouts in univariate analysis, such as differential expression, using mixture modelling [7, 8] but approaches for multivariate problems, including dimensionality reduction, have not yet been considered. As a consequence, it has not been possible to fully ascertain the ramifications of applying dimensionality reduction techniques, such as principal components analysis (PCA), to zero-inflated data.

**Figure 1:**
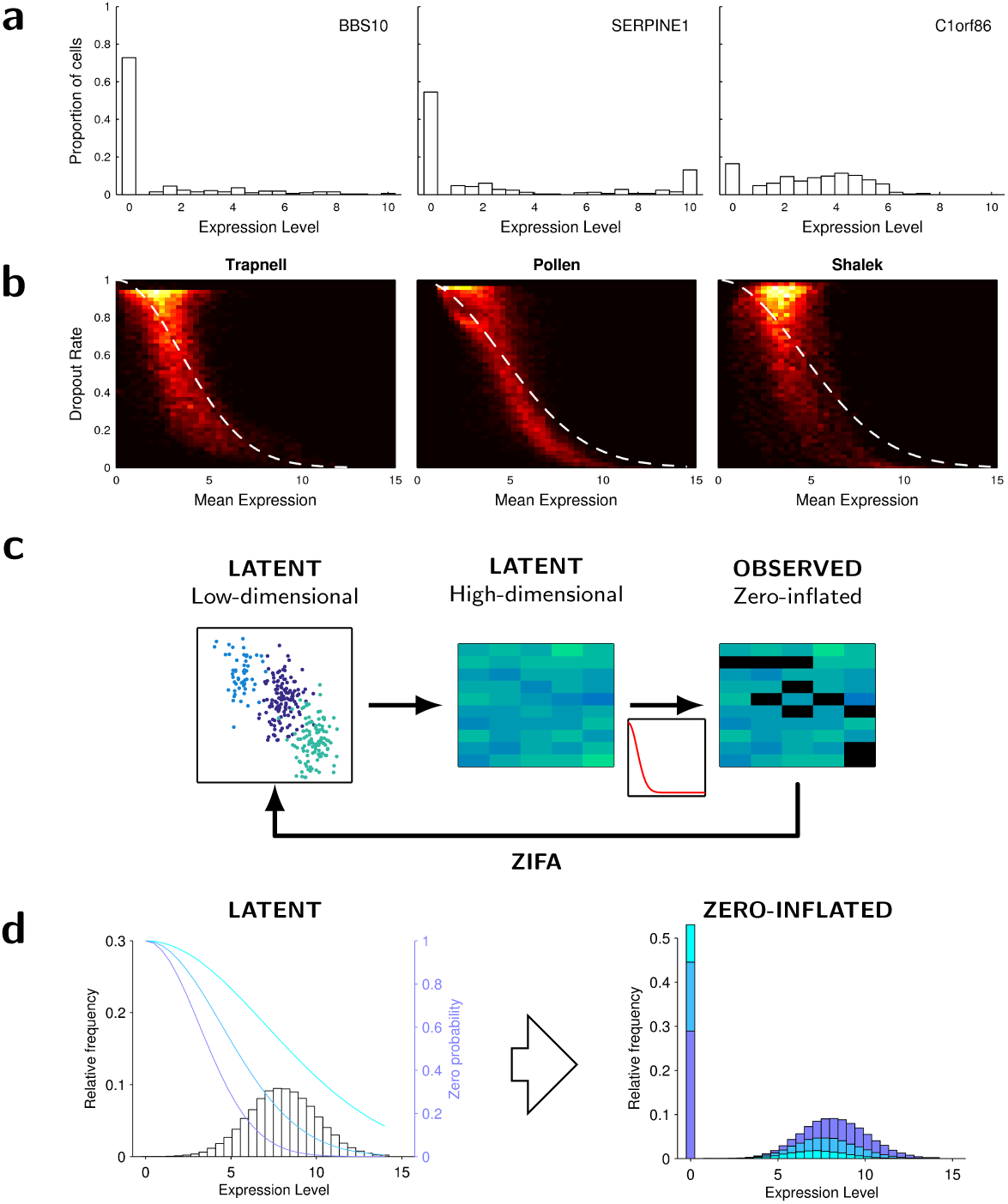
Zero-inflation in single cell expression data. (a) Illustrative distribution of expression levels for three randomly chosen genes shows an abundance of single cells exhibiting null expression [15]. (b) Heatmaps showing the relationship between dropout rate and mean non-zero expression level for three published single cell data sets [3, 5, 13] including an approximate double exponential model fit. (c) Flow diagram illustrating the data generative process used by ZIFA. (d) Illustrative plot showing how different values of *λ* in the dropout-mean expression relationship (blue lines) can modulate the latent gene expression distribution to give a range of observed zero-inflated data.

Dimensionality reduction is a universal data processing step in gene expression analysis. It involves projecting data points from the very high-dimensional gene expression measurement space to a low dimensional *latent* space reducing the analytical problem from a simultaneous examination of 10,000s of individual genes to a much smaller number of (weighted) collections that exploit gene co-expression patterns. In the low dimensional latent space, it is hoped that patterns or connections between data points that are hard/impossible to identify in the high-dimensional space will be easy to visualize.

The most frequently used technique is principal components analysis which identifies the directions of largest variance (principal components) and uses a linear transformation of the data into a latent space spanned by these principal components. The transformation is linear as the coordinates of the data points in the low-dimensional latent space are a weighted sum of the coordinates in the original high-dimensional space with no non-linear transformations used. Other linear techniques include Factor Analysis (FA) which is similar to PCA but focuses on modelling correlations rather than covariances. Many non-linear dimensionality techniques are also available but linear methods are often used in an initial step in any dimensionality reduction processing since non-linear techniques are typically more computationally complex and do not scale well to simultaneously handling many thousands of genes and samples.

In this article we focus on the impact of dropout events on the output of dimensionality reduction algorithms (principally linear approaches) and propose a novel extension of the framework of Probabilistic Principal Components Analysis (PPCA) [9] or Factor Analysis (FA) to account for these events. We show that the performance of standard dimensionality-reduction algorithms on high-dimensional, single cell expression data can be perturbed by the presence of zero-inflation making them sub-optimal. We present a new dimensionality-reduction model, **Z**ero-**I**nflated **F**actor **A**nalysis (ZIFA), to explicitly account for the presence of dropouts. We demonstrate that ZIFA outperforms other methods on simulated data and single cell data from recent scRNA-seq studies.

The fundamental empirical observation that underlies the zero-inflation model in ZIFA is that the dropout rate for a gene depends on the expected expression level of that gene in the population. Genes with lower expression magnitude are more likely to be affected by dropout than genes that are expressed with greater magnitude. In particular, if the mean level of non-zero expression is given by *μ* and the dropout rate for that gene by *p*_0_, we have found that this dropout relationship can be approximately modelled with a parametric form *p*_0_ = exp(−*λμ*^2^), where *λ* is a fitted parameter, based on a double exponential function. This relationship is consistent with previous investigations [7] and holds in many existing single cell datasets (Figure 1b). The use of this parametric form permits fast, tractable linear algebra computations in ZIFA enabling its use on realistically sized datasets in a multivariate setting.

## Method

### Overview

ZIFA adopts a latent variable model based on the Factor Analysis (FA) framework and augments it with an additional zero-inflation modulation layer. Like FA, the data generation process assumes that the separable cell states or sub-types initially exist as points in a latent (unobserved) low-dimensional space. These are then projected onto points in a latent high-dimensional gene expression space via a linear transformation and the addition of Gaussian-distributed measurement noise. Each measurement then has some probability of being set to zero via the dropout model that modulates the latent distribution of expression values. This allows us to account for observed zero-inflated single cell gene expression data (Figure 1c). The scaling parameter in the dropout model can allow for a large range of dropout-expression profiles (Figure 1d).

In the following we provide a more detailed mathematical treatment of the proposed zero-inflated factor analysis model although we leave a complete exposition for the Supplementary Information.

### Statistical Model

Let *N* be the number of samples, *D* be the number of genes, and *K* be the desired number of latent dimensions. The data is given by a high-dimensional *N* × *D* data matrix **Y** = [**y**_1_,…, **y**_*N*_], where *y*_*ij*_ is the level of expression of the *j*-th gene in the *i*-th sample. The data is assumed to be generated from a projection of a latent low-dimensional *N* × *K* matrix **Z** = [**z**_1_,…, **z**_*N*_] (*K* ≪ *D*). In all derivations below, we use use *i* = 1,…, *N* to index over samples (cells), *j* = 1,…, *D* to index over genes, and *k* = 1,…, *K* to index over latent dimensions. Each sample **y**_*i*_ is drawn independently:

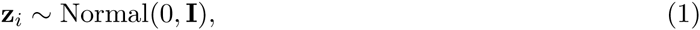

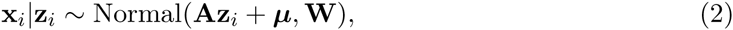

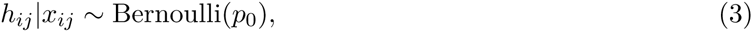

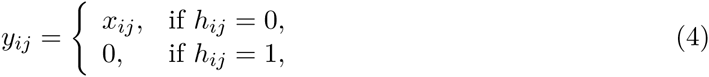

where **I** denotes the *K* × *K* identity matrix, **A** denotes a *D* × *K* factor loadings matrix, **H** is a *D* × *N* masking matrix, 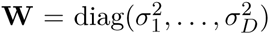 is a *D* × *D* diagonal matrix and ***μ*** is a *D* × 1 mean vector. We choose the drop out probability to be a function of the latent expression level, 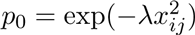, where *λ* is the exponential decay parameter in the zero-inflation model. Note that *λ* is shared across genes which reduces the number of parameters to be estimated and captures the fact that technical noise should have similar effects across genes.

### Statistical Inference

Given an observed single cell gene expression matrix **Y** we wish to identify model parameters Θ = (*A, σ*^2^, *μ, λ*) that maximize the likelihood *p*(**Y***|θ*). We do this using the expectation-maximization (EM) algorithm. We summarize the algorithm in the box below and then describe the algebraic details:

#### Algorithm 1

EM for Zero-Inflated Dimensionality Reduction

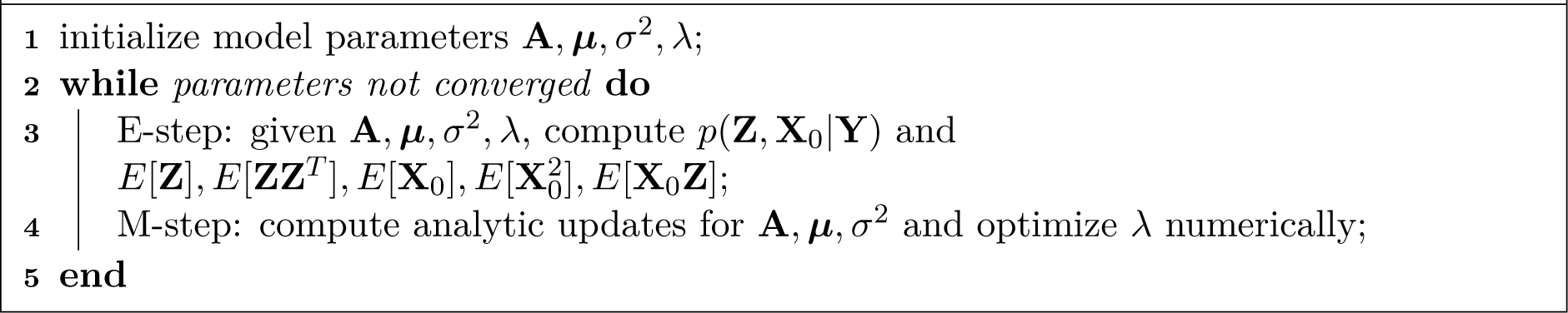

We denote the value of the parameters at the *n*-th iteration, Θ_*n*_, as the value that maximizes the expected value of the complete log likelihood *p*(**Z**, **X**, **H**, **Y**) under the conditional distribution over the latent variables given the observed data and the parameters at the last iteration. Computing the value of the parameters at each iteration requires two steps: the expectation step (E-step) and the maximization step (M-step). In the E-step, we derive an expression for the complete log likelihood *p*(**Z**, **X**, **H**, **Y** *|*Θ_*n*_) and compute all necessary expectations under the distribution *p*(**Z**, **X**, **H** *|***Y**, Θ_*n*−1_). The approximate zero-inflation model that we adopt admits closed form expressions for the expectations allowing the algorithm to be applied to realistically sized datasets. In the M-step, we maximize the expected value of the complete log likelihood with respect to Θ_*n*_.

The EM algorithm structurally resembles the equivalent algorithm for FA that iterates between imputing the coordinates of the observed data points in the low-dimensional latent space (E-step) and optimizing model parameters (M-step). In ZIFA, the expectation step incorporates a data imputation stage to compute the expected gene expression levels for genes/cells with observed null values. Note, that if the noise measurement variance attributed to each gene is identical, we obtain a zero-inflated version of the Probabilistic PCA algorithm [9] (ZI-PPCA).

## Results

### Simulation study

We tested the relative performance of ZIFA against Principal Components Analysis (PCA), Probabilistic PCA (PPCA) [9], Factor Analysis and, for reference, non-linear techniques including Stochastic Neighbour Embedding (t-SNE) [10], Isomap [11], and Multidimensional Scaling (MDS) [12]. First, we generated simulated datasets according to the PPCA/FA data generative model with the addition of one of three dropout models (i) a double exponential model (as assumed by ZIFA), (ii) a linear decay model and (iii) a missing-at-random uniform model. The latter two models were designed to test the robustness of ZIFA to extreme misspecification of the dropout model. Data was simulated under a range of different conditions by varying noise levels, dropout rates, number of latent dimensions and number of genes. The simulation experiment was not intended to truly reflect actual real world data characteristics but to establish, when all other modelling assumptions are true, the impact of dropout events on the outcomes of (P)PCA and FA.

### Setup

We used the assumed generative model to produce simulated data. For the simulations, the values *a*_*jk*_ were drawn from a uniform distribution *U* (−0.5, 0.5), the diagonal elements of the covariance matrix were drawn from a uniform distribution *U*(0.9, 1.1)*σ*^2^, where *σ*^2^ is a simulation parameter, and *μ*_*j*_ were drawn from *U*(2.7, 3.3). We experimented with three choices of *f*(⋅): a decaying squared exponential, 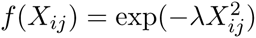 (used in ZIFA); a linear decay function, *f*(*X*_*ij*_) = 1 −*λX*_*ij*_; and a uniform (missing at random) function for each gene *j*, *f*(*X*_*ij*_) = 1 −*λ*_*j*_.

We used a base setting of *N* = 150, *K* = 10, *D* = 50, *σ*^2^ = 0.3, *λ* = 0.1 and explored the effects of altering the decay parameter *λ*, the number of latent dimensions *K*, the cluster spread *σ*^2^, the number of observed dimensions *D*, and the number of samples *N*.

### Performance metrics

As a measure of algorithm performance, we compared the true **z**_*i*_ to the 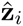 for each sample estimated by the algorithms as follows. We computed the true distance between each pair of points *j, k* and defined a pairwise distance matrix *F* such that *F*_*jk*_ = ‖**z**_*j*_ − **z**_*k*_‖_2_. We compared this to the estimated distance matrix 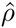 with 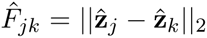. We scored the correspondence between the two distance matrices using Spearman correlation *ρ*_*s*_. By comparing *F* and 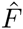 rather than **z**_*i*_ and 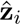,we account for the fact that dimensionality reduction algorithms may rotate the points but ought to preserve the relative distances between them.

### Outcomes

Although the data sets was generated according to a PPCA/FA model (up to the dropout stage), in the presence of cells with genes possessing zero expression, the performance of all standard dimensionality methods (even PPCA/FA) deteriorated relative to ZIFA. Our simulation results (Figure 2b) indicate that standard approaches maybe safely used in certain regimes but should be avoided in others. In particular, gene sets with a high degree of zero-inflation will be problematic (small *λ*), as the relative distances between data points in the gene expression measurement space will be distorted by the presence of zeros and hence there will be a error when projecting back into the latent space. Performance also falls if the gene set is small since there is less scope to exploit strong co-expression signatures across genes to mitigate for the presence of zeros. These regimes are important to consider in the context of linear transformation techniques (PCA, PPCA and FA) that are often applied only to curated gene sets where the linearity constraints maybe approximately applicable. The application of non-linear techniques did not cure the problems induced by dropouts.

**Figure 2:**
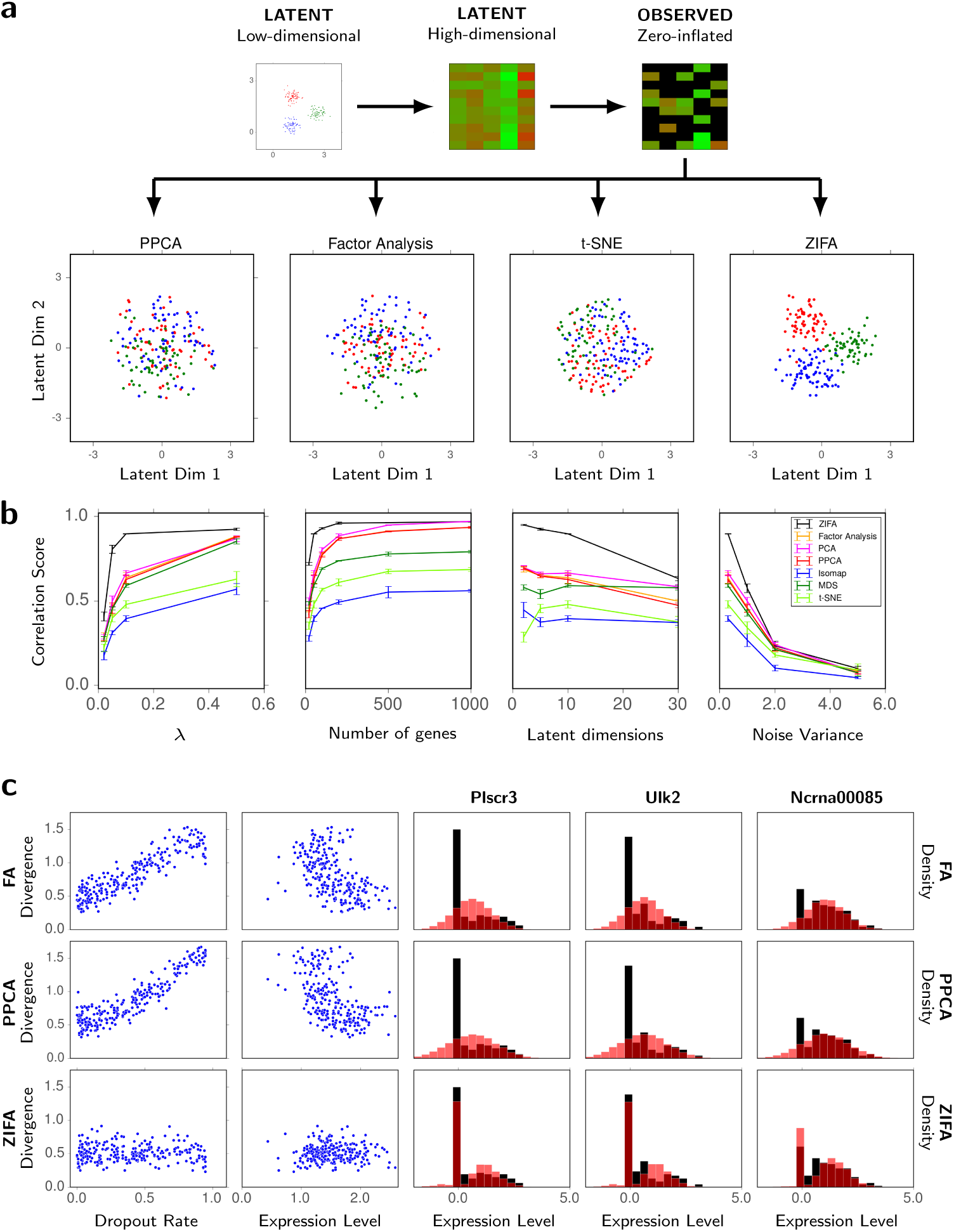
Performance comparison of dimensionality reduction techniques. (a) Toy simulated data example illustrating the performance of ZIFA compared to standard dimensionality reduction algorithms. (b) Performance on simulated datasets based on correlation score between the estimated and true latent distances as a function of *λ* (larger *λ*, lower dropout rate), number of genes and latent dimensions and noise level used in the simulations. (c) Plots showing the divergence between the predictive and empirical data distributions as a function of dropout rate and mean expression level for FA, PPCA and ZIFA. Illustrative predictive performance and model fits (red) on the T-cell single cell data set [3] (black).

Overall, ZIFA outperformed the standard dimensionality reduction algorithms. This would be expected for those simulations adopting the same generative model assumed by ZIFA (Figure 2b) but performance was also replicated regardless of whether dropouts were added following a linear model (Supplementary Fig. 1A), or a missing-at-random model (Supplementary Fig. 1B). This suggests that it is better to account for dropouts somehow even if the dropout characteristics are not realistic. Interestingly, this may suggest that ZIFA could be applicable for other zero-inflated multivariate data sets.

ZIFA should therefore be considered a safe alternative in that it converges in performance to PPCA/FA in the large data, low-noise limit but is robust to dropout events that might distort the outcomes of these methods in non-ideal situations.

### Real data analysis

We next sought to test these methods in an experiment based on real single cell expression datasets [3, 5, 6, 13]. In this case, the “true” latent space is unknown and we are unable to measure performance as with the previous simulated data experiment. Instead, for each of the data sets, we took random subsets of 25, 100, 250 and 1,000 genes and applied ZIFA, PPCA and FA to each subset assuming 5 latent dimensions.

For each gene *j*, we compared the posterior predictive distribution 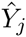 of the distribution of read counts from each method to the observed distribution *Y*_*j*_ as follows: (1) we computed the proportion of values in *Y*_*j*_ and 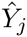 that fell into 30 discrete intervals, (2) we then computed the difference between the histograms Δ_*j*_. If *h*_*n*_ is the proportion of values in bin *n* for the true distribution, and 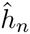 for the predicted distribution, then the histogram divergence is given by

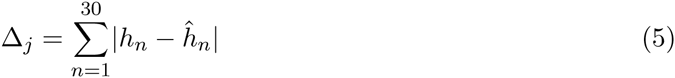

We computed the fraction of genes for which the Δ_*j*_ from ZIFA was less than Δ_*j*_ from PPCA and factor analysis. To prevent overfitting, we assessed fit on a test set: we fit the model for each dataset on a training set containing 70% of the datapoints, and computed the difference between the histograms on the remaining 30% of datapoints.

Note, it is not possible to do this comparison with standard PCA or other dimensionality methods, such as t-SNE, since these are not based on a probabilistic generative model framework and therefore it is not possible to derive the posterior predictive distributions that we use for performance comparisons.

Using this criterion we found that predictive distributions from PPCA and FA showed high divergence for genes that exhibited a high dropout rate or possessed a low non-zero expression level. This meant that the predictive data distributions were a poor fit for the empirical data. ZIFA performance was largely unaffected in contrast (Figure 2c). Example predictive model fits are shown for the T-cell data set [3] for three genes: `Plscr3`, `Ulk2` and `Ncrna00085` (Figure 2c).

The statistical frameworks underlying PPCA and FA employ Gaussianity assumptions that are unable to explicitly account for zero-inflation in single cell expression data. The dropout model used by ZIFA modulates this Gaussianity assumption allowing for zero-inflation leading to drastically improved modelling accuracy. Across the four data sets we found that the predictive distribution derived by ZIFA was superior to those of PPCA and FA on at least 80% of the genes examined and often over 95% (Table 1).

**Table 1:** Comparison of ZIFA to PPCA and FA on four biological datasets. Columns are the number of genes in the dataset (selected at random). Percentages denote the proportion of genes for which ZIFA provided a better fit than FA/PPCA, averaged across 100 replicates.

| Dataset | Method \| | Subset Size |  |  |  |
| --- | --- | --- | --- | --- | --- |
|  |  | 25 | 100 | 250 | 1000 |
| Differentiating T-Cells | FA | 86 ± 6.6% | 84 ± 4.6% | 82 ± 4.9% | 84 ± 8.8% |
| | PPCA | 88 ± 6.3% | 87 ± 4.1% | 89 ± 4.5% | 100 ± 0.3% |
| 11 Populations | FA | 97 ± 3.7% | 96 ± 2.5% | 96 ± 2.1% | 95 ± 2.7% |
| | PPCA | 98 ± 3.2% | 97 ± 2.0% | 97 ± 1.5% | 99 ± 0.6% |
| Myoblasts | FA | 97 ± 3.3% | 97 ± 2.4% | 96 ± 2.7% | 95 ± 2.7% |
| | PPCA | 97 ± 3.2% | 96 ± 2.3% | 96 ± 2.1% | 99 ± 1.7% |
| Bone Marrow | FA | 98 ± 3.0% | 97 ± 2.0% | 97 ± 1.7% | 97 ± 1.7% |
| | PPCA | 98 ± 3.1% | 97 ± 1.8% | 97 ± 1.4% | 97 ± 1.3% |

## Discussion

The density of dropout events in scRNA-seq data can render classical dimensionality-reduction algorithms unsuitable and to-date it has not been possible to assess the potential ramifications of applying such methods on zero-inflated data. We have modified the PPCA/FA framework to account for dropout to a produce a “safe” method for dimensionality reduction of single-cell gene expression data that provides robustness against such uncertainties. In the absence of dropout events, the method is essentially equivalent to PPCA/FA, and therefore software implementations can straightforwardly substitute our approach for existing methods (e.g. `Z = PCA(Y, k)` to `Z = ZIFA(Y, k)`). Our methodology differs from approaches, such as Robust PCA, that model corrupted observations. ZIFA treats dropouts as real observations, not outliers, whose occurrence properties have been characterised using an empirically informed statistical model.

ZIFA models strictly zero measurements rather than near-zero values. It has been possible to account for near-zero values in a univariate mixture modelling framework by placing a small-variance distribution around zero rather than a point mass [7, 8]. Achieving the same goal, in a multivariate context, requires further methodological thought and development in order to produce a solution that is computationally tractable with a large number of dimensions.

Finally, the ZIFA framework lies strictly in the linear transformation framework but non-linear dimensionality reduction approaches, such as t-SNE [10], have proven to be highly effective in single cell expression analysis. It is an area of on-going investigation to determine how zero-inflation can be formally accounted for with such methods. A natural direction would be to directly incorporate it in a non-linear generative approach such as the Gaussian Process Latent Variable Model (GP-LVM) [14]. ZIFA is also potentially applicable to other zero-inflated data where there is a negative correlation between the frequency with which a measurement feature is zero and its mean signal magnitude in non-zero samples.

A Python-based software implementation is available online: https://github.com/epierson9/ZIFA.

## Author’s contributions

E.P. and C.Y. conceived the study and developed the algorithms. E.P. performed data analysis and developed the software implementation. E.P. and C. Y. wrote the manuscript.

## Acknowledgements

E.P. acknowledges support from the Rhodes Trust. C.Y. is supported by a UK Medical Research Council New Investigator Research Grant (Ref. No. MR/L001411/1), the Wellcome Trust Core Award Grant Number 090532/Z/09/Z, the John Fell Oxford University Press (OUP) Research Fund and the Li Ka Shing Foundation via a Oxford-Stanford Big Data in Human Health Seed Grant.

## Additional Files

### Additional file 1 — Supplementary Information

Supplementary Figures and Methods.

